# Low Temperature and High Hydrostatic Pressure Have Compounding Negative Effects on Marine Microbial Motility

**DOI:** 10.1101/2022.10.26.513967

**Authors:** Kelli K. Mullane, Masayoshi Nishiyama, Tatsuo Kurihara, Douglas H. Bartlett

## Abstract

Approximately three fourths of all pelagic marine prokaryotes live in the deep-sea, an environment characterized by low temperature and high hydrostatic pressure. Within deep-sea environments labile organic matter is often scarce and motility can serve as a competitive advantage for microorganisms. Experimental work with a handful of species suggests motility is one of the most temperature- and pressure-sensitive cellular processes, however the combined effects of temperature and pressure together have yet to be investigated in detail. Here we employed growth-dependent motility agar assays and growth-independent microscopy assays to assess how changes in these two physical factors impact motility both individually and in combination, using ecologically relevant model organisms from the cosmopolitan genera *Halomonas, Alcanivorax*, and *Marinobacter*. At pressures equivalent to bathyal and abyssal depths, changes in temperature from 30°C to 4°C (motility assays) or 23°C to 7°C (microscopy assays) had a greater influence on motility than pressure. In addition, low-temperature and high-pressure impacts were additive. Exposure to high pressure had varying degrees of effect on flagellar function, depending on the strain and the magnitude of the pressure. These ranged from short-term impacts that were quickly reversible to long-term impacts that were detrimental to the function of the flagellum, leading to complete loss of motility. These findings highlight the sensitivity of deep-sea bacterial motility systems to combined temperature/pressure conditions, phenotypes that will contribute to the modulation of diverse microbial activities at depth.

**IMPORTANCE:** Microorganisms perform critical functions in biogeochemical cycles at depth, as well as likely modulating the carbon sequestration potential of the deep ocean. However, their activities under in situ conditions are poorly constrained. One aspect of microbial activity is motility, generally mediated by the energy-consuming rotation of one or more flagellar filaments that enables swimming behavior. This provides a competitive advantage for microbes in the environment, such as by enhancing nutrient acquisition. Here we report on culture-based and microscopy-based analyses of pressure-temperature (P-T) effects on the motility of three ecologically relevant marine microbes. The results in all cases indicate that high pressure and low temperature exert compounding inhibitory effects. This argues for the need for further investigations into P-T effects on deep-sea microbial processes.

## INTRODUCTION

The ocean - particularly at the microscale - is remarkably heterogeneous, and marine microorganisms face a wide range of physical and chemical gradients (1). To thrive in such a dynamic environment, microbes utilize motility and chemotaxis to search out favorable niches (2–4). For motile microbes, this microenvironment is largely defined by that organism’s motility range (1); the more motile a microbe (i.e., the faster their swimming speed and the more chemotactic they are), the larger the volume they are able to exploit. Additionally, motility can enhance a microbe’s ability to degrade organic matter by both increasing the rate at which a microbe encounters organic matter (5, 6) and aiding in attachment to particulate organic matter (POM) (3, 7–9). A significant fraction of marine prokaryotes are found in the deep-sea (>1000 meters below sea level, mbsl). Deep-sea pelagic prokaryotes account for nearly 75% of the total prokaryotic biomass and 50% of the total prokaryotic production within the global ocean (10), and abundance estimates for benthic deep-sea environments are comparable to those for seawater (11, 12). In the deep sea, where labile organic matter is less abundant (13–15), microbial motility may be particularly important (6, 16–19). However, these environments are broadly characterized by low temperatures (between 0-4°C) and high hydrostatic pressures (38 Megapascals (MPa) on average) (20), two physical parameters known to impact a wide range of cellular processes, including motility (21, 22). Because marine microorganisms play a pivotal role in global biogeochemical cycling and ocean productivity (23, 24), having a detailed understanding of pressure and temperature (P-T) impacts on cellular processes, including motility, is of fundamental importance, including their consequences to assessments of deep ocean carbon sequestration (25, 26).

Studies of mesophilic microbes indicate that bacterial motility is one of the most temperature-sensitive (27–29) and pressure-sensitive (21, 22, 30, 31) cellular processes. Temperature changes of just a few degrees can dramatically change swimming speeds (28), and pressures as low as 10 MPa (equivalent to ~1,000 mbsl) have been shown to impact both the formation of newly synthesized flagella and the function of preexisting flagella (30). Shallow-water protists may have similar levels of pressure-sensitive motility (32), whereas it is unknown how pressure affects motility within the domain Archaea. Since lower temperatures decrease pressure tolerance in many marine and terrestrial bacterial species (33–41) and decreasing temperature and increasing pressure have additive effects on membrane physical structure (42–45), both parameters should be viewed in concert with one another. Only a handful of studies have assessed the impact of high hydrostatic pressure on microbial motility (27, 30, 31, 34, 46–49). Previous work has assessed the combined influence of low temperature and high hydrostatic pressure on microbial flagellar rotation of the mesophile *Escherichia coli* (27), but to our knowledge, the combined influence of low temperature and high pressure on marine microbial motility is unknown.

In this study, we employ both culture- and microscopy-based analyses to examine the effects of low temperature and high hydrostatic pressure on the motility of three marine bacteria belonging to the genera *Halomonas, Alcanivorax*, and *Shewanella*. These genera are widely distributed within the global ocean, inhabiting a wide range of environments throughout the water column (50–53). This cosmopolitan nature makes them ecologically relevant model organisms. The strains utilized in this study were isolated from the Gulf of Mexico at water depths ranging from 46 to 1509 mbsl, with corresponding *in situ* temperatures ranging from ~23-30°C (54) to ~4-7°C (55). We show that low temperature (4-7°C) has a greater impact than bathyal and abyssal pressures (10-50 MPa), but that the impacts of low temperature and high hydrostatic pressure are compounding.

## MATERIALS AND METHODS

### Study strains

The strains used in this study (Table 1) were generously provided by Dr. Romy Chakraborty and Dr. Gary Andersen at the Lawrence Berkeley National Laboratory. They were isolated from the Gulf of Mexico following the *Deepwater Horizon* oil spill using hydrocarbon-soaked bead traps (56). Briefly, Bio-Traps^®^ (Molecular Insights, Knoxville, TN, USA) were baited with MC-252 crude oil and deployed at multiple depths on a drilling platform approximately 600 meters from the *Deepwater Horizon* platform. The traps were deployed from August through September 2010, and isolates were subsequently obtained using media that contained MC-252 as the sole carbon source. Three strains - 10BA GOM-1509m, 18 GOM-1509m, and 36 GOM-46m (herein referred to as strains 10BA, 18, and 36, respectively) - were chosen for this study. Strains 10BA and 18 were isolated from 1509 mbsl, while strain 36 was isolated from 46 mbsl.

**Table 1:** Summary of study strains utilized in this study, including closest cultured relative (based off 16S rRNA sequencing), isolation depth (meters), optimum growth temperature (°C), optimum growth pressure (Megapascals, MPa), and flagellation type.

| Strain Name | Closest Cultured Relative | Isolation Depth (meters) | Optimum Growth Temperature (°C) | Optimum Growth Pressure (MPa) | Flagellation |
| --- | --- | --- | --- | --- | --- |
| 10BA GOM-1509m | *Halomonas titanicae* | 1509 | 30 | 10 | Peritrichous |
| 18 GOM-1509m | *Alcanivorax xenomutans* | 1509 | 30 | 0.1 | Single Polar |
| 36 GOM-46m | *Shewanella indica* | 46 | 30 | 0.1-10 | Single Polar |

The 16S rRNA gene sequences of the study strains were determined following PCR using the primers 27F and 1492R (57). To obtain near full-length 16S rRNA sequences, Sanger sequencing was performed (Retrogen, Inc., San Diego, CA) with each primer and the sequences were aligned using MEGA X version 10.2.6 (58). These partial 16S sequences were deposited into GenBank with the accession numbers MZ424455 for strain 10BA GOM-1509m, MZ424456 for strain 18 GOM-1509m, and MZ424457 for strain 36 GOM-46m. MEGA X was used to align these sequences with similar 16S rRNA sequences from GenBank, and to construct a maximum-likelihood phylogenetic tree (Figure 1). The flagellation of each strain (Table 1) was determined using a NanoOrange flagellar staining method (59) using a Nikon C1 confocal fluorescence microscope (Nikon, Tokyo, Japan).

**Figure 1.**
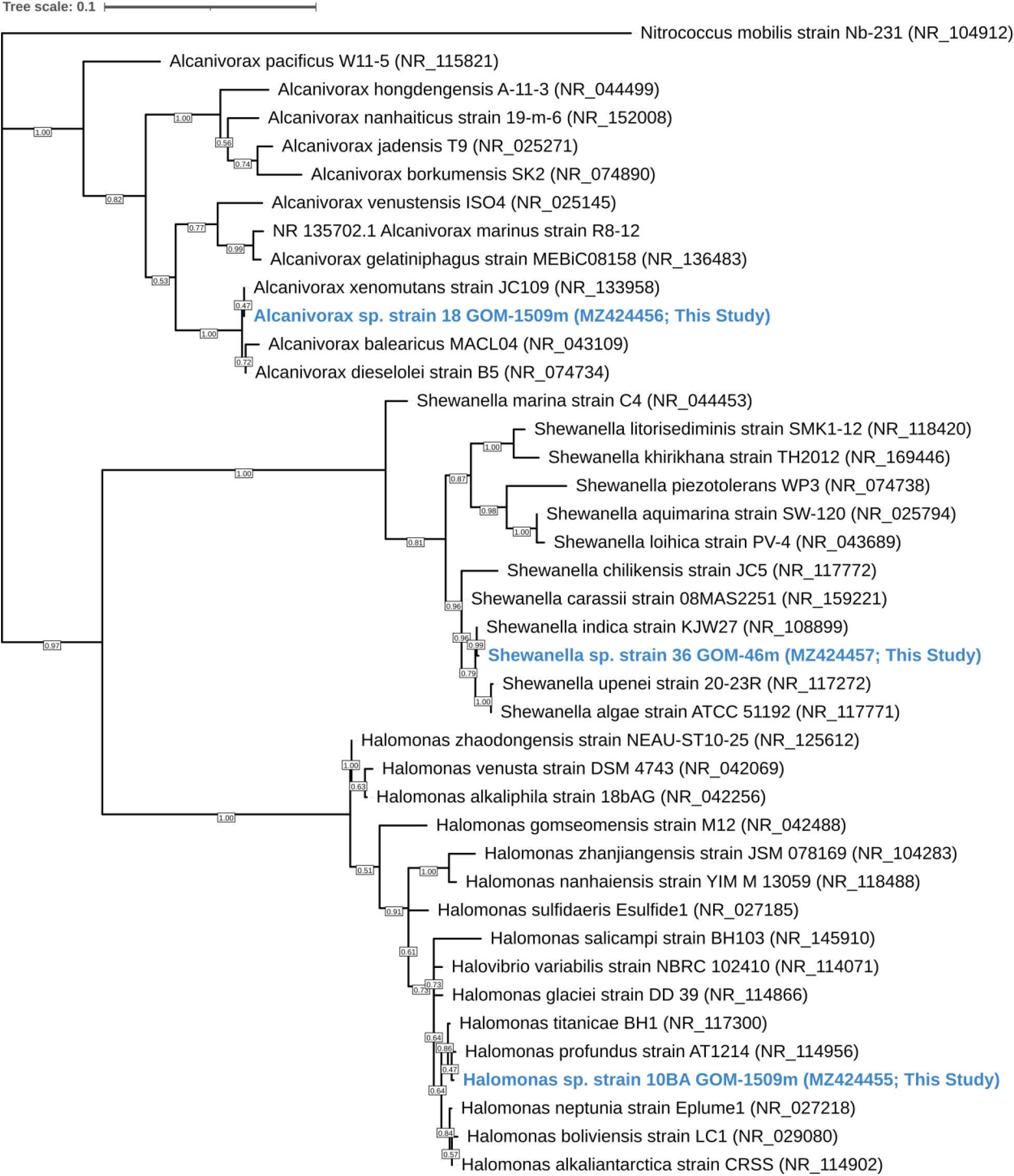
Maximum-likelihood phylogenetic tree based on 16S rRNA gene sequences showing the relationship between strains 10BA GOM-1509m, 18 GOM-1509m, and 36 GOM-46m (blue) and members of the genera *Halomonas, Alcanivorax*, and *Shewanella*, respectively. GenBank accession numbers used are shown in parentheses. The sequence of *Nitrococcus mobilis* strain Nb-231 (NR_104912.1) was used as an outgroup. Bootstrap values (based on 1000 replications) greater than 50% are shown at branch points. Bar, 0.01 substitutions per nucleotide position.

### Growth conditions

All three strains were grown in Marine Broth 2216 (HiMedia Laboratories, Mumbai, India) at a concentration of 28 g/L. When the strains were grown at high hydrostatic pressure the medium was supplemented with a biological buffer (100 mM HEPEs buffer, pH 7.5) and an alternate electron acceptor (100 mM NaNO_3_).

### Pressure and temperature effects on growth

Growth curves were generated at various pressures (0.1, 10, 25, and 50 MPa) and temperatures (4°C and 30°C) to assess the impact of these physical parameters on growth of the three study strains. Briefly, strains were grown to late log phase at the experimental temperature, diluted 1:100 into fresh medium, and transferred into pressurizable polyethylene transfer pipettes. These transfer pipettes were then heat-sealed and incubated in pressure vessels under the appropriate temperature and pressure conditions, as previously described (60). At each timepoint, three transfer pipettes were pulled from each pressure/temperature condition and the optical density at 600nm wavelength (OD_600_) was measured using a GENESYS 10S UV-Vis Spectrophotometer (Thermo Fisher Scientific, Waltham, WA, USA).

### Long-term pressure and temperature effects on motility

To evaluate the effects of pressure and temperature on motility of the study strains, two approaches were employed. The first method was an adaptation of that used by Eloe *et al*. (31), in which a 5 mL glass serum vial (DWK Life Sciences, New Jersey, USA; at 30°C: Catalog No. 223685; at 4°C: Catalog No. 223738) fitted with a pressurizable 2-leg rubber septum (DWK Life Sciences, New Jersey, USA) was filled completely with low percentage (0.3%) agar Marine Medium 2216 supplemented with 100 mM HEPES buffer (pH 7.5) and 100 mM NaNO_3_. Vials were utilized instead of the standard transfer pipette bulbs used for growth assays because vials allowed for significantly improved visualization of microbial motility. We found no statistically significant difference between growth in pressurizable polyethylene pipette bulbs and glass vials fitted with a pressurizable rubber septum (data not shown). Slightly different glass vials were used at 4°C because those used at 30°C tended to break under low temperature, high pressure conditions. Stationary phase cell cultures were inoculated in a straight line through the center of the agar using a needle and syringe. Triplicate vials were incubated under various pressure (0.1, 10, 25, and 50 MPa) and temperature (4°C and 30°C) conditions. Periodically, the vials were briefly removed from the pressure vessels and microbial motility was assessed by measuring visible growth away from the central inoculation site. When determining the distance travelled, the widest part of the cell plume was measured. Tufts of increased motility (see Results) were not included in these measurements.

### Short-term pressure and temperature effects on motility

The second motility evaluation method utilized a high-pressure chamber for optical microscopy, described previously (46). The high-pressure chamber is made of nickel alloy (Hastelloy C276) and is equipped with two optical windows (BK7). It is optimized to obtain high quality images while maintaining a stable hydrostatic pressure up to 150 MPa. The chamber was connected to a hand pump (HP-150, Syn-Corporation, Japan) via a separator, which was used to reduce the total dead volume of buffer solution in the pressure line. The internal temperature of the chamber was controlled by running temperature-regulated water from a thermostat bath through tubing within the chamber. The high-pressure chamber was attached to an inverted microscope (Eclipse T*i*, Nikon, Japan), and microscopic observation was performed by a long-working distance objective lens (CFI ELWD ADM40XC, Nikon, Japan). Phase-contrast videos were acquired by a charge-coupled device camera (WAT-120N+, Watec, Japan).

These experiments were performed at both 23°C and 7°C, which were the closest temperatures the instrument could achieve to the optimum and *in situ* temperatures (30°C and 4°C, respectively) of the strains. The instrument was not capable of reaching temperatures above room temperature and exhibited diminished video quality below 7°C. For these experiments, the strains were grown to mid-log phase in Marine Broth 2216 at the experimental temperature. Cultures were diluted in fresh medium (1:10 or 1:20, depending on cell density), and then gently pipetted into and enclosed within the chamber. The hydrostatic pressure within the chamber was increased to ~100 MPa in increments of 20 MPa, and then decreased in a similar stepwise fashion. Phase-contrast videos of the cells were recorded at 30 frames s^−1^ for 20 seconds at each pressure. The pressure was regulated with an accuracy of ±1 MPa; the experimental temperature was regulated with an accuracy of ±1°C. The total elapsed time for pressure treatment of a population of cells was approximately 4 minutes. After pressure was released, cells were removed from the chamber and the assay was repeated with a fresh culture. All strains were examined in triplicate using biological replicates.

Microscopy data was analyzed using previously established methods (61). Briefly, the percentage of motile cells was determined by selecting a 1 second portion of the video file and manually counting the motile and non-motile cells in the plane of focus. A total of 10 frames per video were analyzed, giving an n = 30 across all 3 replicates for each pressure. For swimming speed, the SimplePTA plug-in (61) for Fiji (62) was used to track cells and determine the cell’s average velocity. A total of 10 cells per video were tracked, giving an n = 30 across all 3 replicates for each pressure. Relatively large standard deviations are common with this type of assay, as has been previously (46).

### Recovery of swimming motility after pressure exposure

The recovery of swimming motility after pressure treatment was also assessed using the high-pressure microscope system. Microbial cultures were grown to mid log phase in Marine Broth 2216 and inserted into the chamber as described above. After recording a 20 second video to document motility of the strain prior to high hydrostatic pressure exposure, the pressure was increased to 80, 120, and 100 MPa for strains 10BA, 18, and 36, respectively. These were the maximum pressures each strain could withstand during the short-term high-pressure microscope experiments. After approximately 3 minutes of exposure to these high pressures, the video camera was turned on and the pressure was decreased to 0.1 MPa, after which recording continued for approximately 3 minutes. These experiments were all performed at 23°C. Using the data analysis methods described above, both the percentage of motile cells and the swimming speed were determined before, during, and after exposure to high hydrostatic pressure.

## RESULTS

### Strain identification and characterization

Strains 10BA, 18, and 36 were isolated from the Gulf of Mexico at water depths of 46, 1509, and 1509 mbsl, respectively. Corresponding *in situ* water temperatures were ~23-30°C at 46 mbsl (54), and ~4.7°C at 1509 mbsl (55). Comparison of the 16S rRNA gene sequence of each study strain with sequences present in GenBank indicated that strain 10BA GOM-1509m belongs to the genus *Halomonas*, strain 18 GOM-1509m to the genus *Alcanivorax*, and strain 36 GOM-46m to the genus *Shewanella*. Based on Maximum-Likelihood phylogenetic analyses (Figure 1) the closest phylogenetic relatives of strain 10BA are *H. titanicae* BH1 (NR_117300) and *H. profundus* strain AT1214 (NR_114956), with 99.63%, and 99.78% 16S rRNA gene sequence similarity, respectively. The closest phylogenetic relatives of strain 18 are *A. xenomutans* strain JC109 (NR_13358), *A. dieselolei* strain B5 (NR_074734), and *A. balearicus* MACL04 (NR_043109), with 100.00%, 99.78%, and 98.90% 16S rRNA gene sequence similarity, respectively. The closest phylogenetic relatives of strain 36 are *S. indica* strain KJW27 (NR_108899), *S. algae* strain ATCC 51192 (NR_117771), and *S. upenei* strain 20-23R (NR_117272), with 99.86%, 99.22%, and 99.07% 16S rRNA gene sequence similarity, respectively.

All three study strains grow optimally in Marine Broth 2216 at 30°C (data not shown). NanoOrange flagellar staining (59) revealed that *Halomonas* sp. strain 10BA is peritrichously flagellated, consistent with what is known about motile members of the genus *Halomonas* (63). The genus *Alcanivorax* contains both motile and non-motile members, and those who are motile have been found to employ a variety of flagellation types (peritrichous, single polar) (64, 65). Our analysis revealed that *Alcanivorax* sp. strain 18 is motile by a single polar flagellum. Finally, *Shewanella* sp. strain 36 is motile by a single polar flagellum, which is consistent with most members of this genus (66), although other flagellation types have been seen in some *Shewanella* species (49, 67).

### Pressure and temperature effects on growth

*Halomonas* sp. strain 10BA exhibited its maximum growth rate and yield when grown at 30°C and 10 MPa (Figure 2A, Supplementary Table S1), making it a piezomesophile, albeit a modest one (68). While slightly diminished growth was observed at 0.1 and 25 MPa, strain 10BA did not grow at 30°C and 50 MPa. At this optimum temperature, its upper pressure limit is comparable to that of the piezotolerant mesophile *Escherichia coli* (33, 69). When grown at 4°C, the pressure optimum of strain 10BA was reduced to 0.1 MPa (Figure 2B, Supplementary Table S1). At this lower temperature, growth rate and yield decreased as pressure increased and no growth was observed at 4°C and 50 MPa.

**Figure 2.**
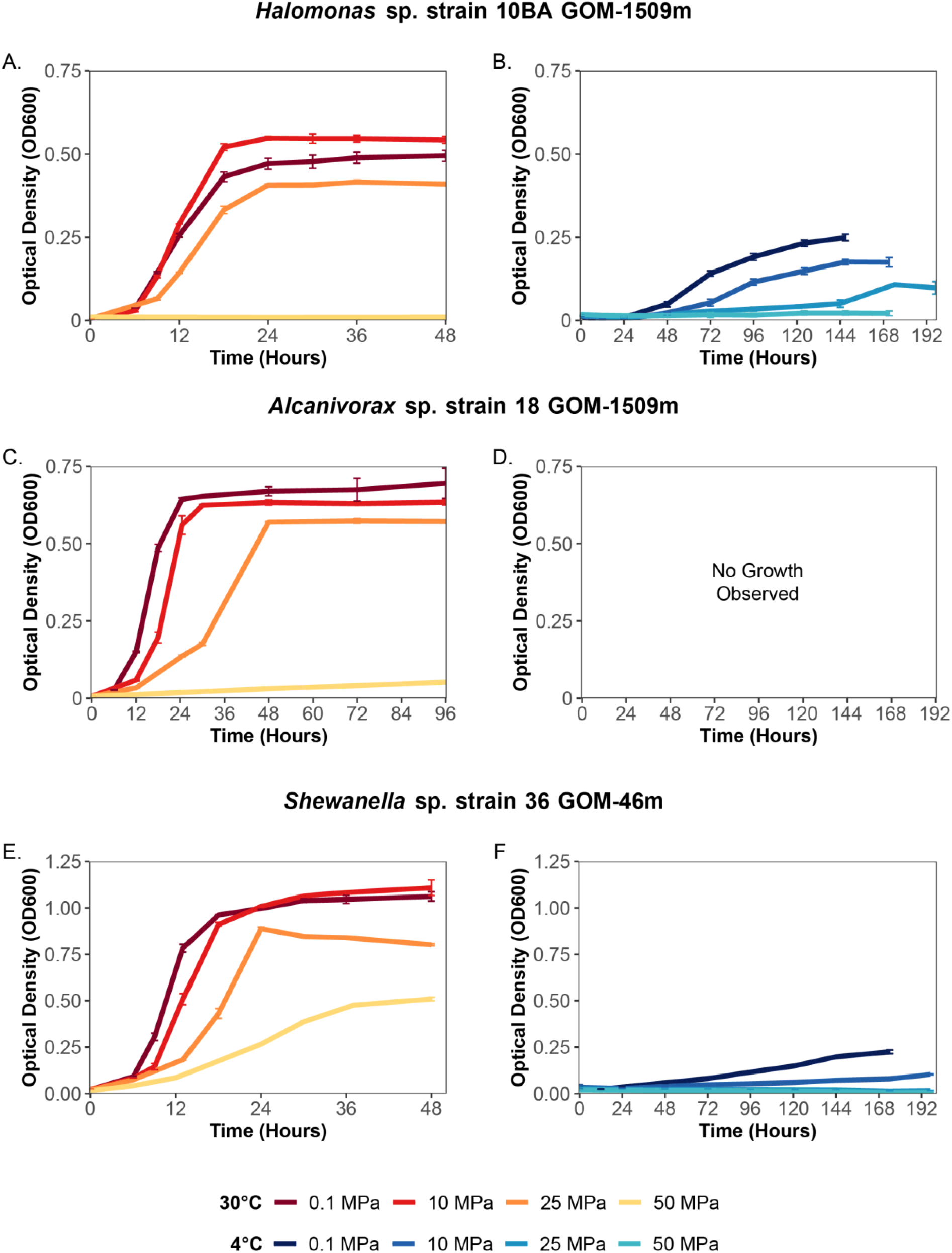
Growth curves of *Halomonas* sp. strain 10BA GOM-1509m at 30°C (A) and 4°C (B), *Alcanivorax* sp. strain 18 GOM-1509m at 30°C (C) and 4°C (D), and *Shewanella* sp. strain 36 GOM-46m at 30°C (E) and 4°C (F) as a function of pressure (0.1, 10, 25, and 50 MPa). Error bars represent standard deviation.

*Alcanivorax* sp. strain 18 displayed more pressure sensitive mesophilic growth properties. At 30°C, *Alcanivorax* sp. strain 18 exhibited its maximum growth rate and yield at atmospheric pressure (Figure 2C, Supplementary Table S1). Growth was only slightly diminished at 10 MPa, and while growth yield only decreased an additional 10% at 25 MPa, the growth rate decreased by over 50%. Growth was barely detectable at 50 MPa for this strain. Strain 18 did not grow at 4°C under the conditions tested.

*Shewanella* sp. strain 36 exhibited similar growth at 0.1 MPa and 10 MPa at 30°C, reaching its maximum growth rate at 0.1 MPa, and its maximum growth yield at 10 MPa (Figure 2D, Supplementary Table S1). An increase in pressure led to a gradual decrease in growth, but unlike the other strains in this study, strain 36 exhibited growth at 30°C and 50 MPa. It is curious that this strain’s growth under its optimum temperature is the most pressure tolerant of the strains despite being isolated near the sea surface at 46 mbsl. However, this piezotolerance was not accompanied by concomitant adaptation to low temperature. When grown at 4°C, the maximum growth rate and yield were at atmospheric pressure (Figure 2E, Supplementary Table S1). At this lower temperature, just 10 MPa worth of pressure led to an approximately 50% decrease in growth rate and yield, and no growth was observed at 25 or 50 MPa.

### Long-term pressure and temperature effects on motility

The effect of pressure and temperature on the growth and motility of the strains was evaluated using pressurizable serum vials containing low percentage motility agar media. For this assay, we calculated the P_1/2_, which we defined as the pressure at which motility is 50% of the value measured at atmospheric pressure (see Supplementary Methods). In general, the pressure and temperature impacts on the motility of these strains was qualitatively similar to those on growth; the better the cells grew, the more motility they exhibited.

*Halomonas* sp. strain 10BA was motile at 0.1, 10, and 25 MPa at 30°C (Figure 3A) but was only motile at 0.1 and 10 MPa at 4°C (Figure 3B). At 30°C, strain 10BA was slightly more motile at 10 MPa than 0.1 MPa, but this pressure effect was not observed at 4°C (see Supplementary Figure S1). At 25 MPa, motility was greatly diminished, with the swimming rate less than 20% that observed at 10 MPa (Figure 3A). As previously established, strain 10BA did not grow at 50 MPa and 30°C, and thus no growth or motility were observed under these conditions. The P_1/2_ for strain 10BA was 11 MPa at 4°C and 23 MPa at 30°C (Supplementary Table S2).

**Figure 3.**
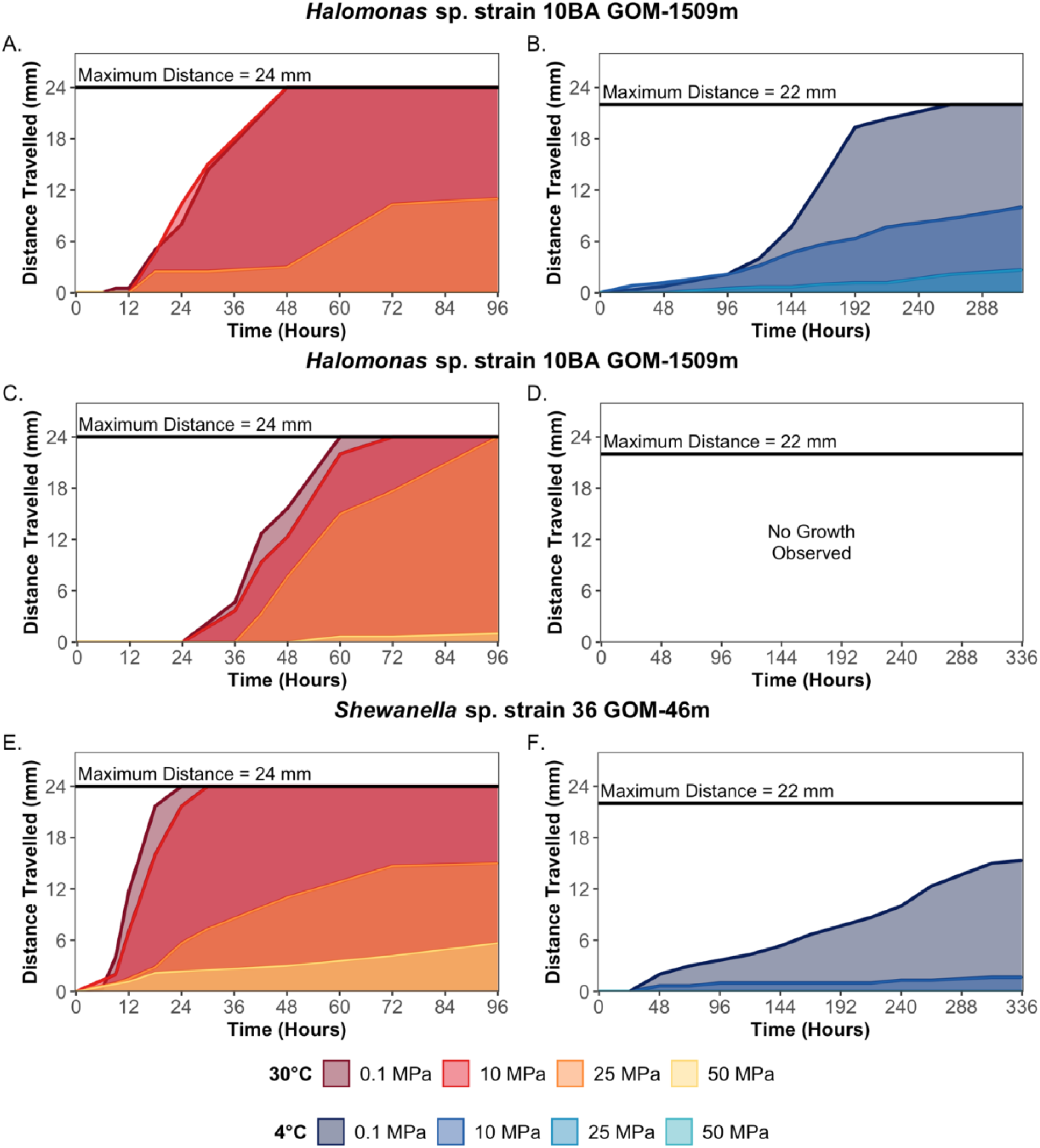
Long-term motility of *Halomonas* sp. strain 10BA GOM-1509m at 30°C (A) and 4°C (B), *Alcanivorax* sp. strain 18 GOM-1509m at 30°C (C) and 4°C (D), and *Shewanella* sp. strain 36 GOM-46m at 30°C (E) and 4°C (F). Assessed as a function of pressure (0.1, 10, 25, and 50 MPa) using a glass serum vial fitted with a pressurizable septum.

*Alcanivorax* sp. strain 18 was motile at 0.1, 10, and 25 MPa at 30°C (Figure 3C). The motility of this strain was only mildly impacted by pressure; the swimming rate at 25 MPa was approximately 50% that observed at 0.1 MPa. Strain 18 was not motile at 50 MPa but did grow at the point of inoculation (see Supplementary Figure S3). Since strain 18 did not grow at 4°C, no motility vial experiments were performed at this temperature. The P_1/2_ for strain 18 was 27 MPa at 30°C (Supplementary Table S2).

*Shewanella* sp. strain 36 was motile at 0.1, 10, and 25 MPa at 30°C (Figure 3E) but was only motile at 0.1 MPa at 4°C (Figure 3F). At 30°C the motility of this strain was comparable at 0.1 and 10 MPa. At 50 MPa and 30°C, *Shewanella* sp. strain 36 grew near the inoculation point, but was not motile. However, for a single replicate, a small tuft of motile cells derived from the inoculation site was observed at the 96-hour time point (Supplementary Figure S4), possibly the result of a mutation (70). The P_1/2_ for strain 36 is 6 MPa at 4°C and 21 MPa at 30°C (Supplementary Table S2).

### Short-term pressure and temperature effects on motility

A high-pressure microscope system (described in Methods) was used to assess the impacts of pressure and temperature on swimming motility over short periods of time (seconds to minutes). For this assay, we also calculated P_1/2_ values, which in this case were defined as the pressure at which either the percentage of motile cells or the swimming speed is 50% that determined at atmospheric pressure (see Supplementary Methods). A representative compilation of the videos acquired during this experiment can be found in the Supplementary Materials (Supplementary Video 1).

*Halomonas* sp. strain 10BA was pressurized to a maximum pressure of 80 MPa at 23°C and 60 MPa at 7°C (Figure 4A & B). At 23°C, the percentage of motile cells remained steady up to 60 MPa, above which it decreased drastically. Similarly, at 7°C, the percentage of motile cells remained steady up to 40 MPa, above which it decreased (Figure 4A). For both temperatures, the swimming speed decreased at a steady rate throughout the pressurization process (Figure 4B). For strain 10BA, the P_1/2_ for percentage of motile cells was 64 MPa at 23°C and 57 MPa at 7°C; the P_1/2_ for swimming speed was 66 MPa at 23°C and 63 MPa at 7°C (Supplementary Table S3). Upon depressurization to 0.1 MPa at 23°C, the percentage of motile cells was significantly higher (mean ± standard deviation; 80% ± 7%) than at the onset of the experiment (62% ± 9%) (p-value << 0.005) (Figure 4A), and the swimming speed recovered to approximately the same level observed before pressurization (Figure 4B). After depressurization at 7°C, both the percentage of motile cells and swimming speed recovered to starting levels (Figure 4A & B).

**Figure 4.**
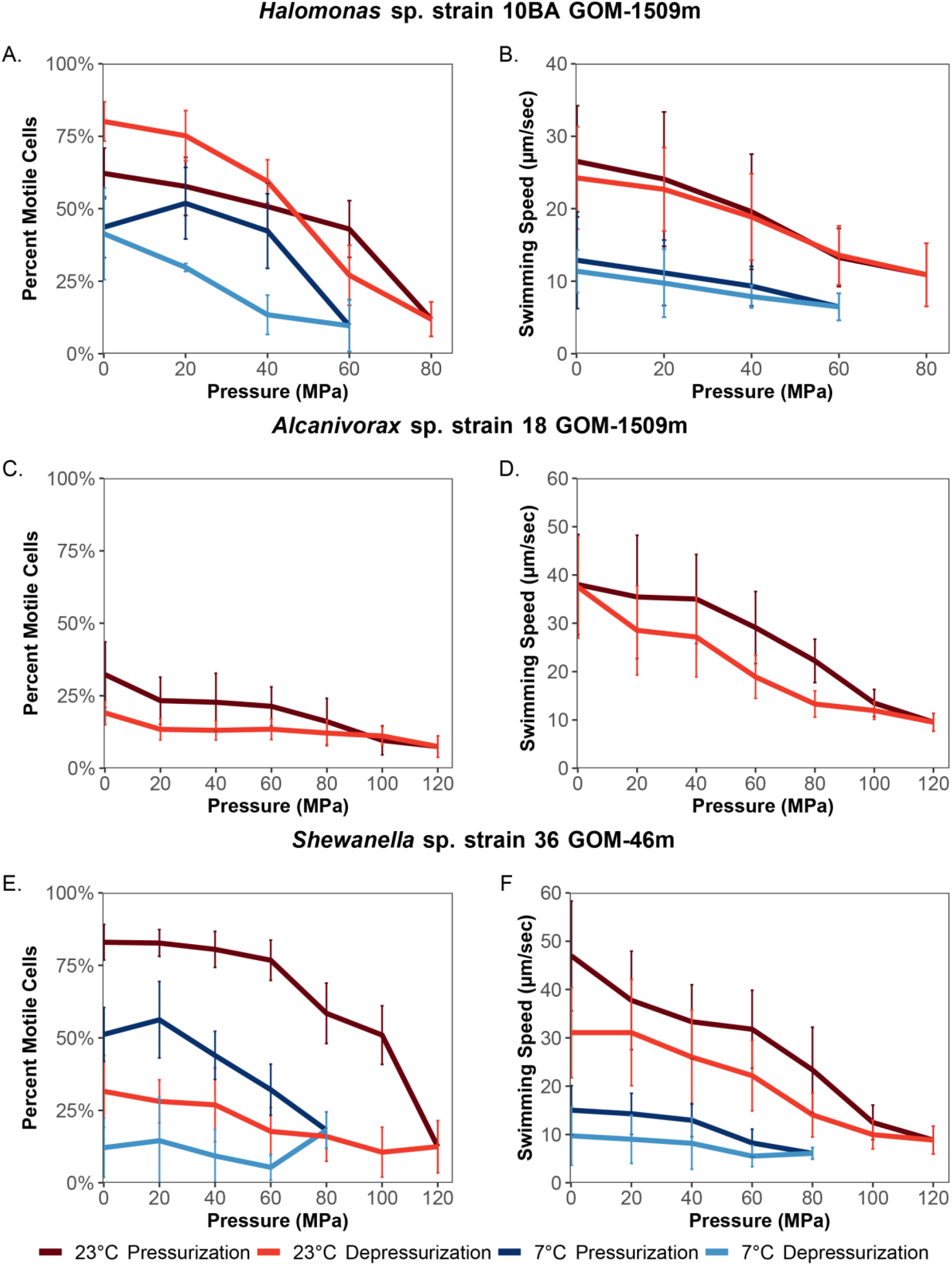
Percentage of motile cells and swimming speed (μm/second) of *Halomonas* sp. strain 10BA GOM-1509m (A & B), *Alcanivorax* sp. strain 18 GOM-1509m (C & D), and *Shewanella* sp. strain 36 GOM-46m (E & F). Assessed as a function of pressure and temperature (23°C and 7°C) using a high-pressure microscope chamber. Error bars represent standard deviation.

*Alcanivorax* sp. strain 18 was pressurized to a maximum pressure of 120 MPa at 23°C (Figure 4C & D). The percentage of motile cells was low at the onset of the experiment in comparison to strain 10BA (32% ± 11% and 62% ± 9%, respectively), and steadily decreased as pressure increased (Figure 4C). The swimming speed remained stable up to 40 MPa, above which it gradually decreased (Figure 4D). The P_1/2_ is 76 MPa for percentage of motile cells, and 88 MPa for swimming speed. Upon depressurization, the swimming speed fully recovered to the level observed at the onset of the experiment (Figure 4D), while the percentage of motile cells only recovered to approximately 60% of the starting value (p-value << 0.005) (Figure 4C).

*Shewanella* sp. strain 36 was pressurized to a maximum pressure of 120 MPa at 23°C and 80 MPa at 7°C (Figure 4E & F). The percentage of motile cells at 23°C decreased in a stepwise fashion as pressure increased. In contrast, at 7°C the percentage of motile cells was relatively stable up to 20 MPa, above which it decreased; the slight increase in percentage of motile cells observed at 20 MPa is not statistically significant (p-value = 0.23) (Figure 4E). At 23°C, the swimming speed gradually decreased as pressure increased. At 7°C, the swimming speed at the onset of the experiment was nearly 70% lower than that observed at 23°C (47 ± 11 μm/s at 23°C, 15 ± 5 μm/s at 7°C). At this lower temperature, the swimming speed remained steady until 40 MPa, above which it decreased more significantly (Figure 4F). The P_1/2_ for percentage of motile cells is 102 MPa at 23°C and 73 MPa at 7°C; the P_1/2_ for swimming speed is 74 MPa at 23°C and 72 MPa at 7°C. Upon depressurization to 0.1 MPa, the percentage of motile cells only recovered to 38% of the value observed prior to pressurization at 23°C, and 24% of that observed prior to pressurization at 7°C (Figure 4E). After depressurization, the swimming speed recovered to 66% of that present prior to pressurization at 23°C, and 65% of that present before pressurization at 7°C. Although the error bars overlap, these represent statistically significant decreases in swimming speed (p-value << 0.005 for 23°C and < 0.005 for 7°C) (Figure 4F).

To rule out cell death caused by exposure to high hydrostatic pressure as an explanation for the reduction in motility in the strains, we evaluated the colony forming units (CFU) before and after pressurization (Supplementary Methods). No significant change in CFU was observed for any of the strains at either 23°C or 7°C (Supplementary Figure S6).

### Recovery of swimming motility after pressure exposure

Since some of the cells exhibit significant loss of swimming motility (both the percentage of motile cells and swimming speed) after depressurization, the recovery of motility was tracked after a 3 minute high pressure exposure at 23°C (Figure 5). Each strain was exposed to the maximum pressure at which it maintained motility during the short-term pressure exposure experiment −80, 120, and 100 MPa for strains 10BA, 18, and 36, respectively.

**Figure 5.**
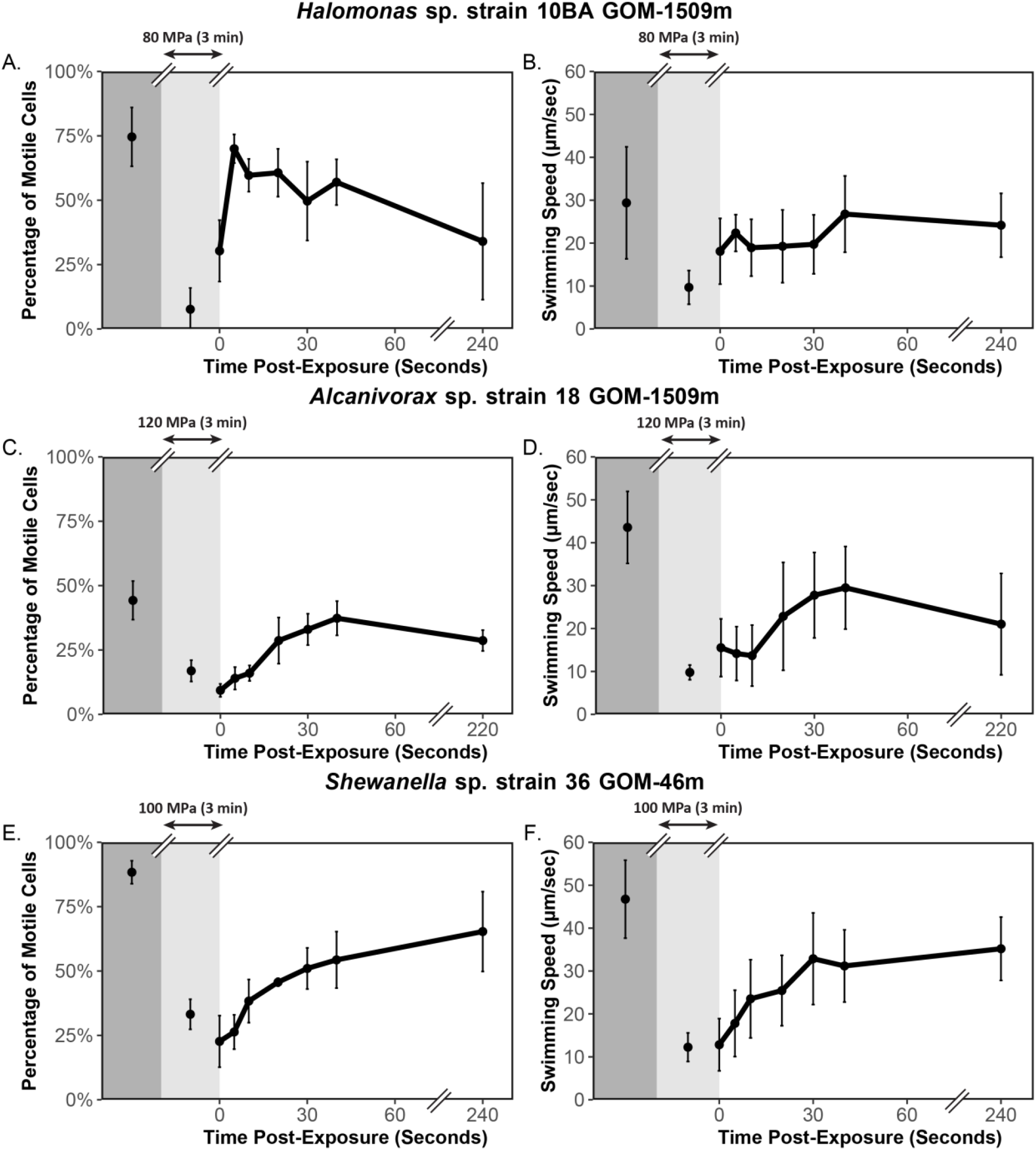
Percentage of motile cells and swimming speed of study strains before (dark gray shading), during (light gray shading), and immediately after (white shading) exposure to pressure at 23°C. *Halomonas* sp. strain 10BA GOM-1509m (A & B) was pressurized to 80 MPa; no data for replicate 3 at 240 seconds post-exposure. *Alcanivorax* sp. strain 18 GOM-1509m (C & D) was pressurized to 120 MPa. *Shewanella* sp. strain 36 GOM-46m (E & F) was pressurized to 100 MPa. Error bars represent standard deviation.

For *Halomonas* sp. strain 10BA, the percentage of motile cells recovered quickly; just 5 seconds after depressurization, the percentage of motile cells was not significantly different from that observed before pressurization (Figure 5A). Likewise, the swimming speed recovered to levels equivalent to those observed pre-exposure within 40 seconds post-exposure (Figure 5B).

For *Alcanivorax* sp. strain 18, the percentage of motile cells required only 40 seconds to recover post-exposure (Figure 5C), while the swimming speed was still significantly different from pre-exposure levels nearly 4 minutes post-exposure (Figure 5D). These results contrast with those from the short-term motility experiments (Figure 4). The observed difference in the percentage of motile cells data could be due to hysteresis. The cells were tracked for less than 30 seconds after decompression during the short-term motility experiments, which may not have been enough time for the cells to regain swimming activity after pressure release. This phenomenon has been observed in other studies assessing the impacts of short-term pressure exposure on motility (46). In the case of the swimming speed results, it appears that longer exposure to high hydrostatic pressure damages some aspect of the energetics or mechanics of flagellar function in this strain.

Finally, for *Shewanella* sp. strain 36, both the percentage of motile cells and the swimming speed were significantly different from pre-exposure levels after more than 4 minutes post-exposure (Figure 5E & F). It should be noted that in both the short-term motility experiments (Figure 4) and the motility recovery experiments (Figure 5), each strain was differentially impacted by exposure to high hydrostatic pressure.

## DISCUSSION

Microbial motility plays an important role within the ocean, and despite the fact that the deepsea is an environment largely defined by low temperature and high hydrostatic pressure, which comprises over 88% of the ocean volume (20), very little is known about the influence of these physical parameters on marine microbial motility. In this study the impact of low temperature and high hydrostatic pressure on microbial motility were assessed both individually and in combination. This was achieved using a modified motility agar assay to look at long-term (growth-dependent) effects, and a high-pressure microscopy system to look at short-term (growth-independent) effects. The representative strains used in this study, belonging to the ecologically relevant genera *Halomonas, Alcanivorax*, and *Marinobacter*, were isolated from the Gulf of Mexico (56). Their isolation depth was a poor indicator of pressure and temperature tolerance under the conditions tested, presumably because the strains were not isolated under *in situ* pressure and temperature. Under almost all experimental conditions evaluated in this study, increased pressure and decreased temperature led to decreased growth and motility.

Major differences were found in the P-T effects on motility when comparing the long-term and short-term motility assays. When incubated at elevated pressure over the course of days to weeks, the P_1/2_ of motility in semi-solid agar was only ~25 MPa and 10 MPa at 30°C and 4°C, respectively (Supplementary Table S2). Conversely, when exposed to pressure for only seconds to minutes in the microscopy assays, the 50% decrease in the percentage of motile cells and the swimming speed respectively required pressures of at least 64 MPa and 66 MPa at 23°C, and 57 MPa and 63 MPa at 7°C (Supplementary Table S3). These stark differences inherently arise from the nature of the two methods. While the short-term, growth-independent motility assay uncovers the immediate impacts of pressure on flagellar function *in vivo*, the long-term, growth-dependent assay requires cells to reproduce and form flagella in addition to requiring the flagella to be functional. Flagellar function alone is energetically costly (19, 71), and flagellar formation is an additional energetic expense (72) that involves the biosynthesis of six different components (73) involving at least 24 core genes, with some bacteria requiring over 50 genes (74, 75).

Generally, decreased temperature (30°C versus 4°C for long-term assays, 23°C versus 7°C for short-term assays) alone had a significantly larger impact on growth and motility than the pressures employed. Furthermore, the combination of low temperature and high hydrostatic pressure exerted compounding deleterious effects on motility. For example, in the long-term motility assay, the swimming rate of *Shewanella* sp. strain 36 under 25 MPa at the optimum growth temperature (30°C) was more than three times that observed under atmospheric pressure (0.1 MPa) at 4°C (see Supplementary Methods and Results). Additionally, it took significantly less pressure to decrease motility by 50% at colder temperatures, as reflected by the lower P_1/2_ values at 4-7°C compared to 23-30°C (Supplementary Tables S2 & S3). This highlights the importance of taking both *in situ* temperature and pressure into consideration when studying microbial processes in the deep ocean, and this surely includes a variety of global biogeochemical cycles (23, 24).

One possible target of P-T effects on microbial cells which could influence motility is the lipid membrane. Previous studies have similarly shown that low temperature and high pressure have compounding negative effects on these structures (22), with changes in the unsaturated fatty acid composition of membranes being one example of a lipid change which is essential for bacterial adaptation in the deep sea (42, 76). It has been estimated that for the phase state of lipid membranes, a change in pressure of 100 MPa is equivalent to a change in temperature of 20°C (22). We estimated that for the growth-based motility assays examined in this study, a change in pressure of only 40 MPa was equivalent to a change in temperature of 20°C (see Supplementary Methods & Supplementary Results), whereas the short-term microscopy assays had values much closer to that observed for lipids.

Although beyond the scope of this study, there are many possible explanations for the additive, deleterious impacts of low temperature and high pressure on motility. First, both high pressure and low temperature are well-known to impact protein synthesis and protein-protein quaternary interactions (77–83), which could impact flagellar synthesis and assembly. In fact, the assembly of flagellar and motility proteins into a large and complex flagellar organelle structures is estimated to be much more sensitive to pressure than synthesis of other protein assembly proteins (30). Second, decreased ion flux through the rotor/stator motor components could impact torque generation, and thus hook and filament rotation, as pressure has been shown to impact similar processes in various bacteria (80, 84–86). Similarly, temperature has been shown to impact the rotational speed of flagellar motors, possibly due to impacts on ion translocation (87, 88).

Some intriguing findings suggest possible future lines of fruitful investigation. First, the motility of each study strain was differentially impacted by short-term exposure to high pressure, suggesting that the motility or flagella of various marine microbes may display large variations in P-T impacts. Assessing whether or not this is indeed the case, as well as the mechanistic underpinnings of these variations, would be valuable. Second, despite being a piezomesophile, *Halomonas* sp. strain 10BA’s motility system is not found to function well at high pressure. This finding contrasts the only other study to assess the impact of pressure on microbial motility in a piezophile (31). In that case, *Photobacterium profundum* SS9’s flagellar systems were found to be extremely well adapted for operation under high hydrostatic pressure. Further motility analyses on a broader range of piezophiles could help to clarify the boundaries that exist between piezophilic growth and high pressure-adapted motility. Finally, evidence of pressure impacts on chemotaxis were observed in one of the study strains. *Alcanivorax* sp. strain 18 appeared to exhibit a “run-reverse-flick” swimming pattern, which has been found in other monotrichously flagellated bacteria as a form of chemotaxis (89–93). In one of the replicates, a clear decrease in the frequency of this “run-reverse-flick” pattern was observed at 23°C when pressure increased from 0.1 to 20 MPa (Supplementary Figure S7; Supplementary Video S2), which could be indicative of a decrease in ability to randomize orientation and hence engage in the biased random walk that is at the heart of chemotaxis. One prior study (27) showed that, at 20°C, extremely high hydrostatic pressure (≥120 MPa) was required to induce a counterclockwise to clockwise reversal of the flagellar motor in an *E. coli* mutant deficient in chemotaxis signaling (Δ*cheY*), while at 9°C, pressures ≤60 MPa were required to induce reversal. Further investigation into P-T influences on not just motility, but biased motion in the direction of nutrient sources is also therefore warranted. The effects of both of these processes, as a function of the rich diversity of prokaryotes present, could have profound consequences on the rates of microbially-mediated chemical transformations at depth.

## DATA AVAILABILITY

Raw data from this study was deposited into the Gulf of Mexico Research Initiative database, GRIIDC (UDI: R5.x285.000:0006, R5.x285.000:0016). Partial 16S rRNA gene sequences for the study strains used in this paper can be found in GenBank with the accession numbers MZ424455, MZ424456, and MZ424457.

## AUTHORS’ CONTRIBUTIONS

KKM and MN performed experimental work. KKM, MN, TK, and DHB performed data analysis. KKM and DHB wrote the manuscript. All authors read and approved the final manuscript.

## FUNDING

This work was supported by the Gulf of Mexico Research Initiative for the RFP V Investigator Grant entitled “Role of Microbial Motility for Degradation of Dispersed Oil” (GOMRI 2015-V-328) (GOMRI R-16-0037) to D.B., and by JSPS KAKENHI (JP15H01319 to M.N., JP17H04598 to T.K.).

## ACKNOWLEDGEMENTS

We would like to thank Dr. Romy Chakraborty, Dr. Gary Andersen, and Dr. Stephen Techtmann for generously providing the microbial strains used as part of this study. We would also like to thank Dr. Jun Kawamoto and Dr. Takuya Ogawa for arranging the microscope study. Finally, we would like to thank Dr. Alvaro Muñoz-Plominsky, Luke Fisher, and Miguel Desmarais for reading, commenting on, and enhancing drafts of this manuscript.

## SUPPLEMENTARY METHODS

### Cell viability after pressure exposure

To rule out cell death as a cause for observed decreases in percentage of motile cells after exposure to high hydrostatic pressure, we assessed cell viability before and after exposure to high hydrostatic pressure. For this, strains were grown to mid to late log phase in Marine Broth 2216 at both 30°C and 4°C. An aliquot of each culture was taken, diluted serially in 1x phosphate buffered saline (PBS) and spread-plated on 1.7% agar Marine Broth 2216 plates. Cultures were then transferred to pressurizable polyethylene transfer pipettes. The transfer pipettes were heat-sealed and placed into pressure vessels that had been pre-incubated at either 30°C or 4°C. Pressure was increased in increments of 20 MPa, and then decreased in increments of 20 MPa to replicate the short-term pressure experiment conditions. *Halomonas* sp. strain 10BA was pressurized to 80 MPa at 30°C and 60 MPa at 4°C; *Alcanivorax* sp. strain 18 was pressurized to 120 MPa at 30°C; *Shewanella* sp. strain 36 was pressurized to 120 MPa at 30°C and 80 MPa at 4°C. After depressurization, an aliquot was taken from each transfer pipette was taken, diluted serially in 1x PBS, and spread-plated on 1.7% agar Marine Broth 2216 plates. All plates were incubated at 30°C (optimum growth temperature for all study strains) for 24-48 hours, and colony forming units were counted. This was performed in triplicate (biological replicates) for each study strain at each temperature.

### P_1/2_ calculations

To assist in our comparison of pressure and temperature impacts on swimming motility in the long-term and short-term motility assays described above, we calculated the P_1/2_, which we define as the pressure at which motility is 50% that observed at atmospheric pressure. For the longterm motility experiments, we calculated a swimming rate during the exponential phase for each temperature/pressure condition tested. Swimming rate was calculated using the equation:

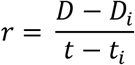

Where *r* = swimming rate, *t_i_* = time at the onset of swimming, *t* = time at the end of swimming (i.e., when the edge of the vial is reached or when the experiment ended if the edge was never reached), *D_i_* = distance travelled at time *t_i_*, and *D* = distance travelled at time *t*. These swimming rates were plotted as a function of pressure and fitted with a linear regression line. The equation for this regression line was then used to determine the pressure at which the swimming rate was 50% that observed at 0.1 MPa.

For the short-term motility experiments, a linear regression line was fitted to the percentage of motile cells vs. pressure and swimming speed vs. pressure graphs (Figure 4). The equation for this regression line was used to determine the pressure at which the percentage of motile cells and the swimming speed were 50% that observed at 0.1 MPa.

### Cell trajectories

To obtain cell trajectories from the high-pressure microscopy videos (Supplementary Figure S7), cells were manually tracked using Fiji (62). Briefly, videos were opened in Fiji and image properties were edited to reflect pixel size and frame interval for the camera used in this study. For each cell being tracked, the curser was hovered over the center of the cell and XY coordinates were recorded. This was repeated every other frame for a total of 1 second of video.

### Pressure-temperature relationship calculations

To obtain a general understanding of the relationship between high pressure and low temperature impacts on growth, growth-dependent motility (vial assay), and growth-independent motility (microscopy assay), ‘rate’ at the differing temperatures were plotted as a function of pressure. For growth, this meant growth rate vs. pressure at 30°C and 4°C; for the growth-dependent motility assay, this meant swimming rate vs. pressure at 30°C and 4°C; and for the growth-independent motility assay, this meant swimming speed and percentage of motile cells vs. pressure at 23°C and 7°C. Growth rate was calculated during the exponential growth phase (Figure 2) using the equation:

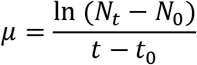

Where *μ* = growth rate constant, *t*_0_ = time at the start of exponential phase, *t* = time at the end of exponential phase, *N*_0_ = optical density at *t*_0_, and *N_t_* = optical density at time *t*. Swimming rate was calculated as described above (see “P_1/2_ calculations” in Supplementary Methods). Linear regression lines were added to these plots for each temperature. Only pressures at which growth was observed were used to generate these regression lines. The x coordinate of the interception of the two linear regression lines was determined, which is a rough estimation of the pressure at which the change in temperature (ΔT) is equivalent to the change in pressure (ΔP). For the growth rate and growth-dependent motility assays, ΔT = 26°C; for the growth-independent motility assays, ΔT = 16°C. Finally, the ΔP/ΔT values were normalized to a ΔT of 20°C.

## SUPPLEMENTARY TABLES

**Supplementary Table S1.** Summary of growth rate (μ, hr^−1^) and maximum optical density at 600nm absorbance (OD600_max_) for study strains when grown at varying temperatures (4°C and 30°C) and pressures (0.1, 10, 25, and 50 MPa).

| Study Strain | Temperature | Pressure (MPa) | Growth Rate ($\mu$, $\text{hr}^{-1}$) | $\text{OD}_{600\text{max}}$ |
| --- | --- | --- | --- | --- |
| *Halomonas* sp. strain 10BA | 4°C | 0.1 | 0.028 | 0.249 |
|  |  | 10 | 0.020 | 0.176 |
|  |  | 25 | 0.018 | 0.108 |
|  |  | 50 | 0.000 | 0.022 |
|  | 30°C | 0.1 | 0.122 | 0.495 |
|  |  | 10 | 0.240 | 0.548 |
|  |  | 25 | 0.181 | 0.416 |
|  |  | 50 | 0.000 | 0.010 |
| *Alcanivorax* sp. strain 18 | 30°C | 0.1 | 0.196 | 0.695 |
|  |  | 10 | 0.161 | 0.643 |
|  |  | 25 | 0.061 | 0.583 |
|  |  | 50 | 0.011 | 0.077 |
| *Shewanella* sp. strain 36 | 4°C | 0.1 | 0.013 | 0.225 |
|  |  | 10 | 0.005 | 0.104 |
|  |  | 25 | 0.000 | 0.028 |
|  |  | 50 | 0.000 | 0.021 |
|  | 30°C | 0.1 | 0.235 | 1.062 |
|  |  | 10 | 0.204 | 1.108 |
|  |  | 25 | 0.144 | 0.888 |
|  |  | 50 | 0.084 | 0.510 |

**Supplementary Table S2.** P_1/2_ values for long-term motility experiments using the glass serum vials at 4°C and 30°C. The P_1/2_ is the pressure at which the swimming rate is half that observed at atmospheric pressure.

| Study Strain | Temperature | $P_{1/2}$ |
| --- | --- | --- |
| *Halomonas* sp. strain 10BA | 4°C | 11 MPa |
|  | 30°C | 23 MPa |
| *Alcanivorax* sp. strain 18 | 30°C | 27 MPa |
| *Shewanella* sp. strain 36 | 4°C | 6 MPa |
|  | 30°C | 21 MPa |

**Supplementary Table S3.**
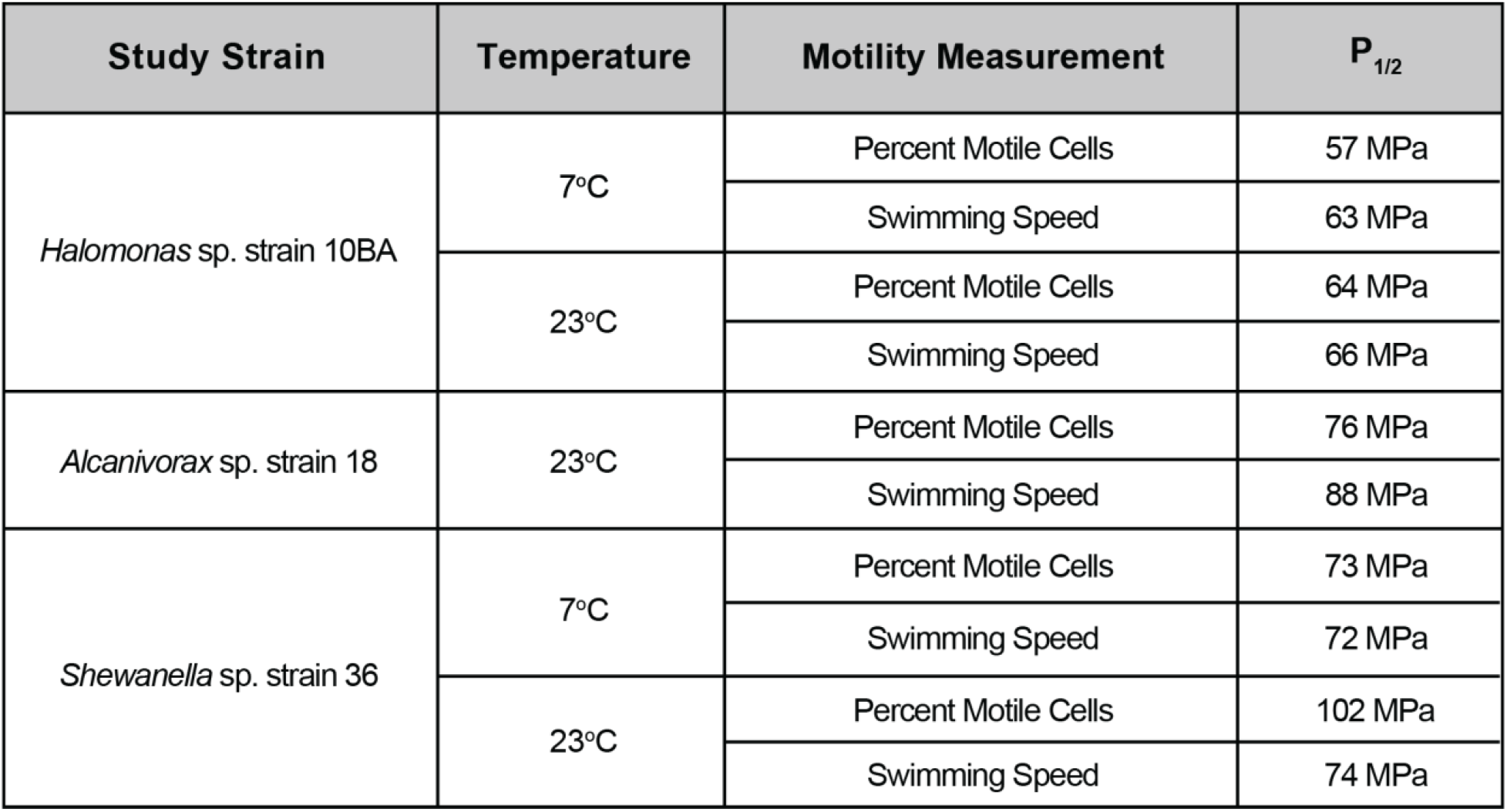
P_1/2_ values for the short-term motility experiments using the high-pressure microscope chamber at 7°C and 23°C. The P_1/2_ is the pressure at which the percentage of motile cells or the swimming speed is half that observed at atmospheric pressure.

## SUPPLEMENTARY FIGURE LEGENDS

**Supplementary Figure S1.**
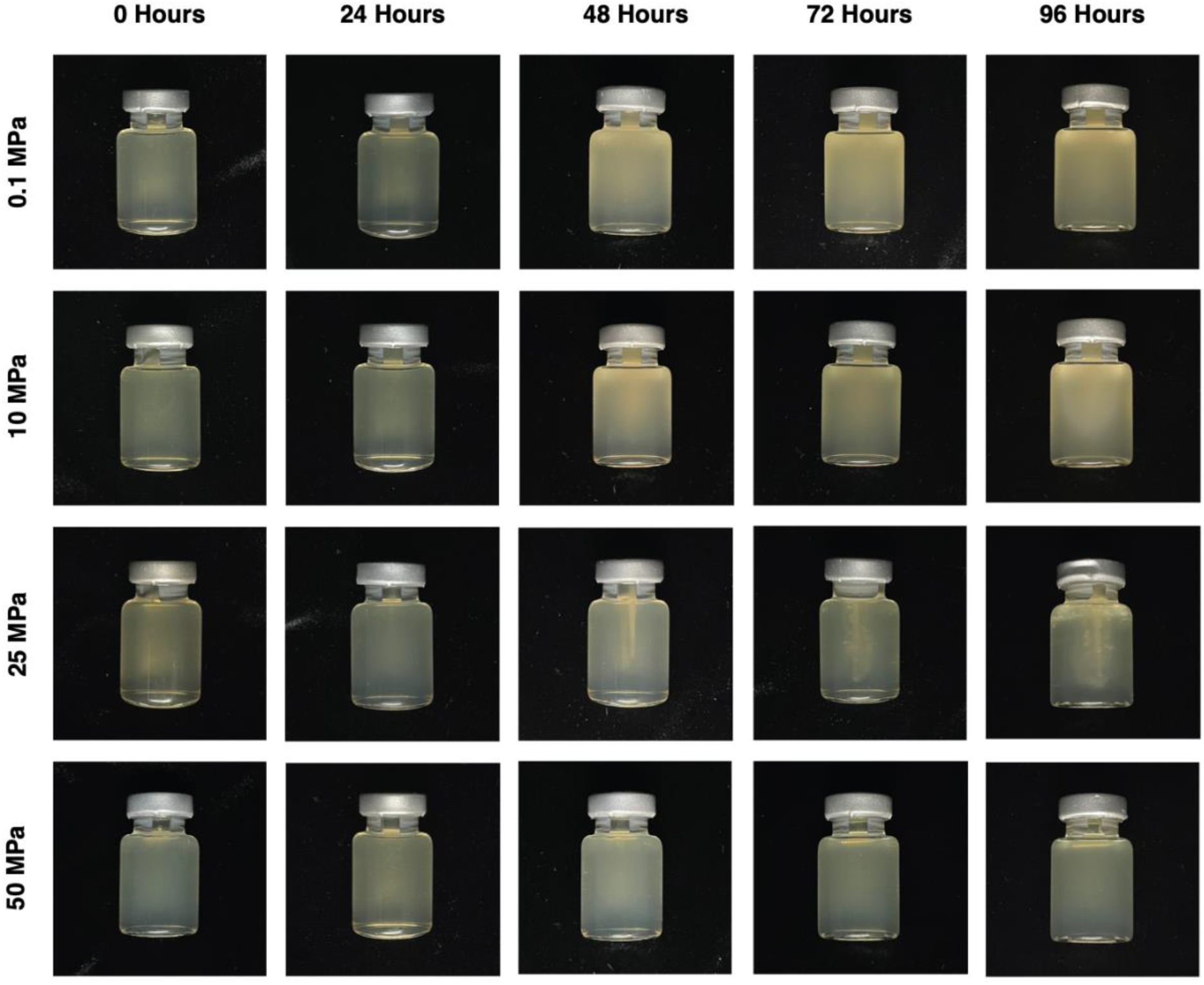
Photos of *Halomonas* sp. strain 10BA GOM-1509m long-term motility vials (30°C) as a function of pressure (y-axis) and time (x-axis).

**Supplementary Figure S2.**
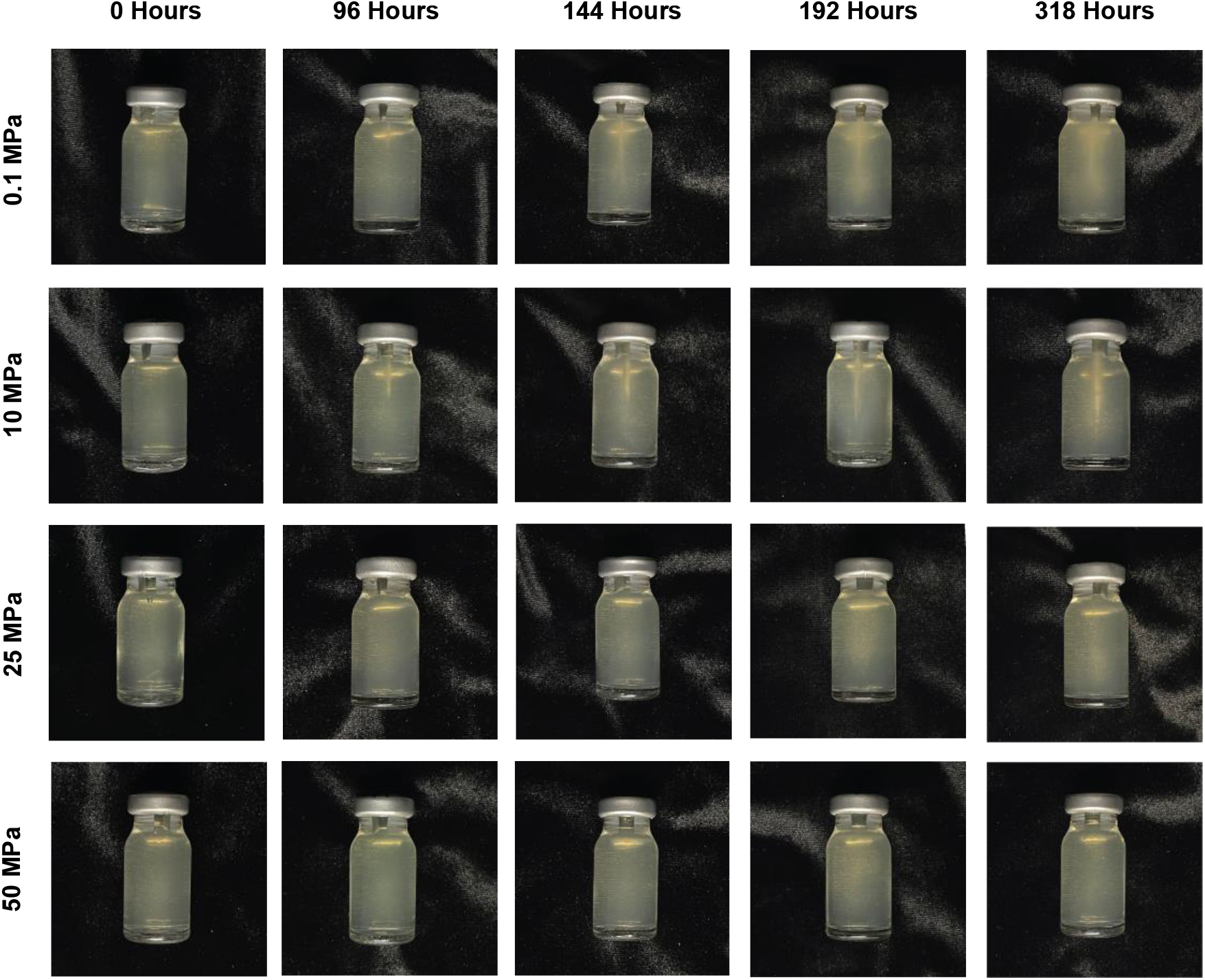
Photos of *Halomonas* sp. strain 10BA GOM-1509m long-term motility vials (4°C) as a function of pressure (y-axis) and time (x-axis).

**Supplementary Figure S3.**
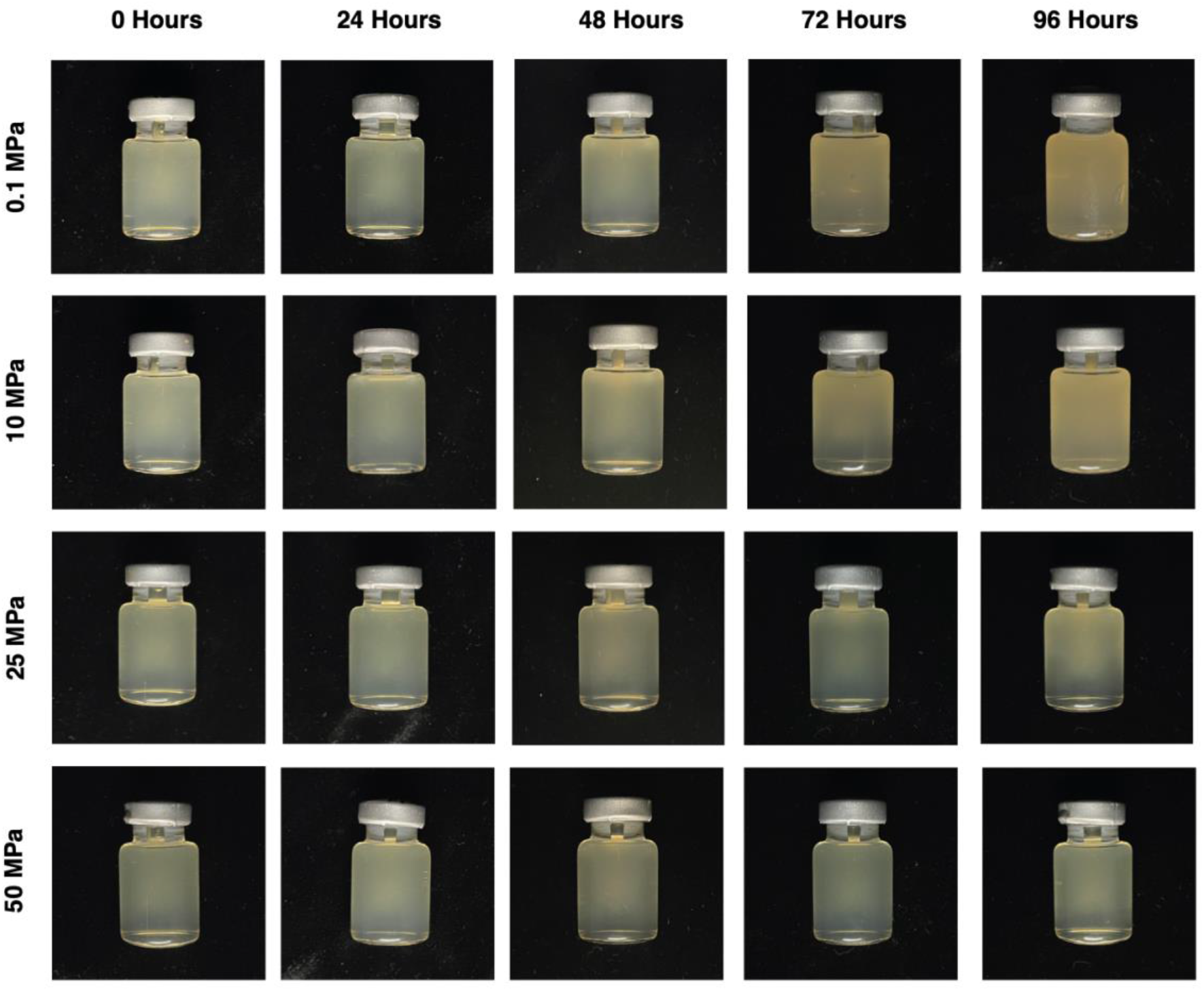
Photos of *Alcanivorax* sp. strain 18 GOM-1509m long-term motility vials (30°C) as a function of pressure (y-axis) and time (x-axis).

**Supplementary Figure S4.**
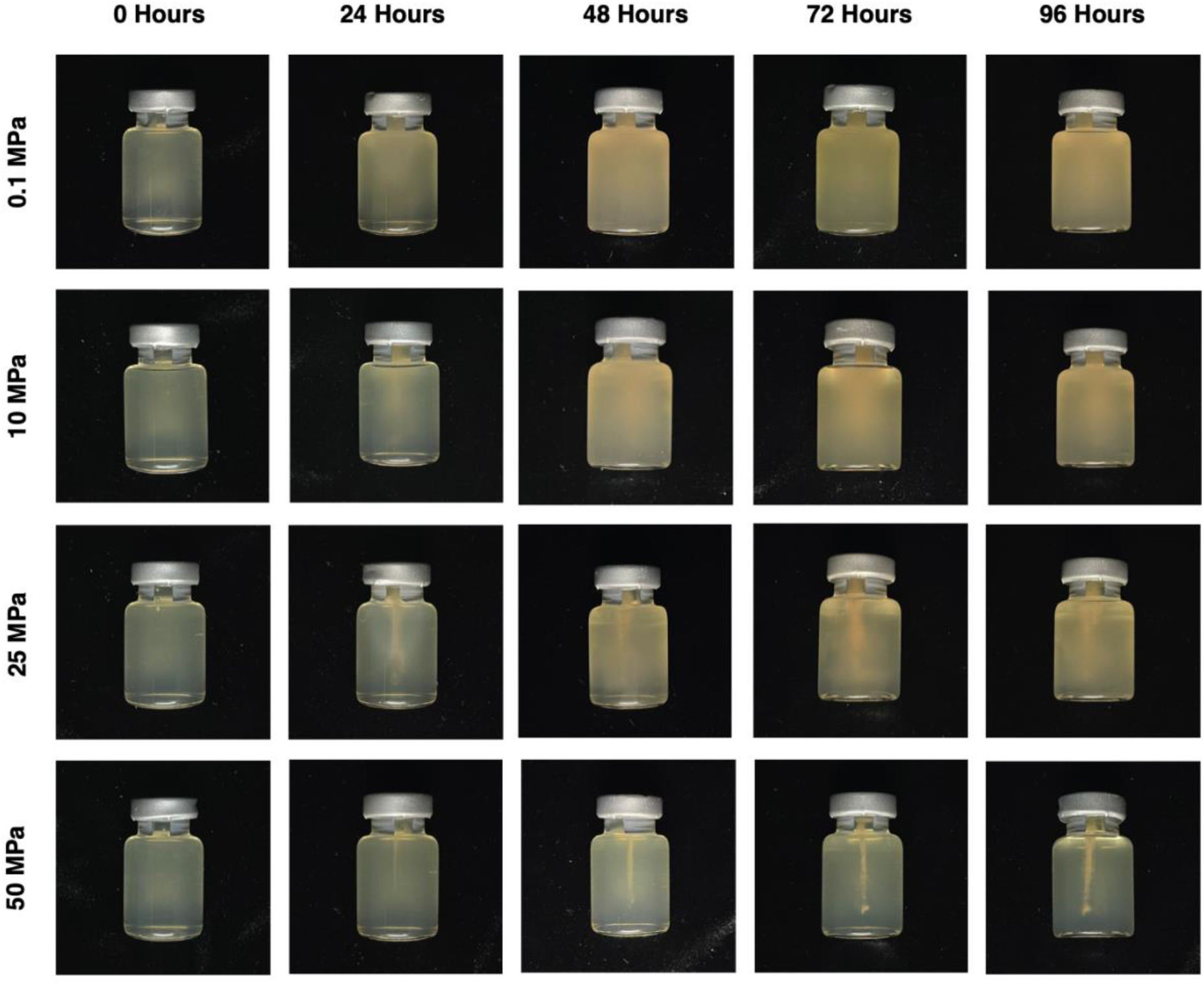
Photos of *Shewanella* sp. strain 36 GOM-46m long-term motility vials (30°C) as a function of pressure (y-axis) and time (x-axis).

**Supplementary Figure S5.**
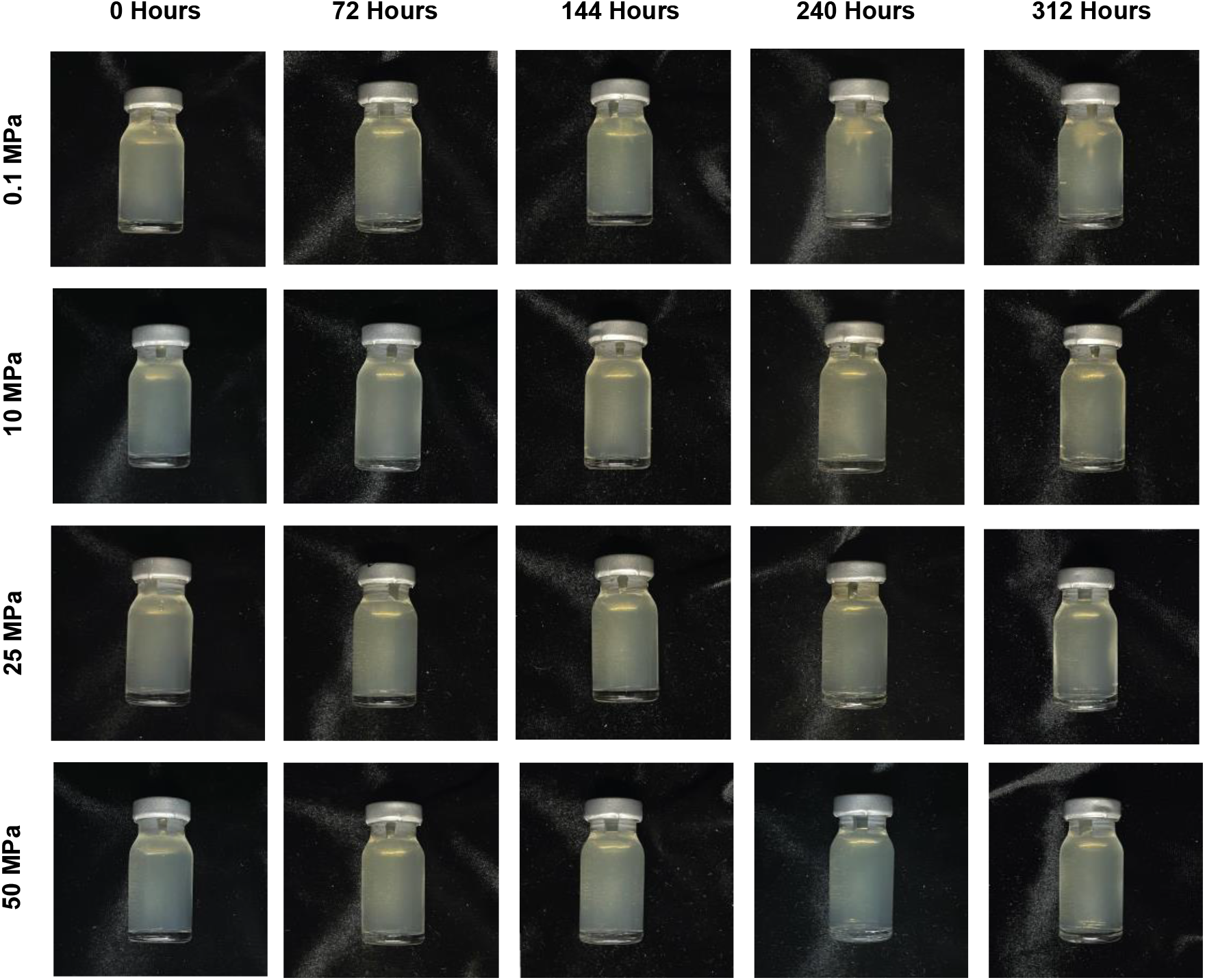
Photos of *Shewanella* sp. strain 36 GOM-46m long-term motility vials (4°C) as a function of pressure (y-axis) and time (x-axis).

**Supplementary Figure S6.**
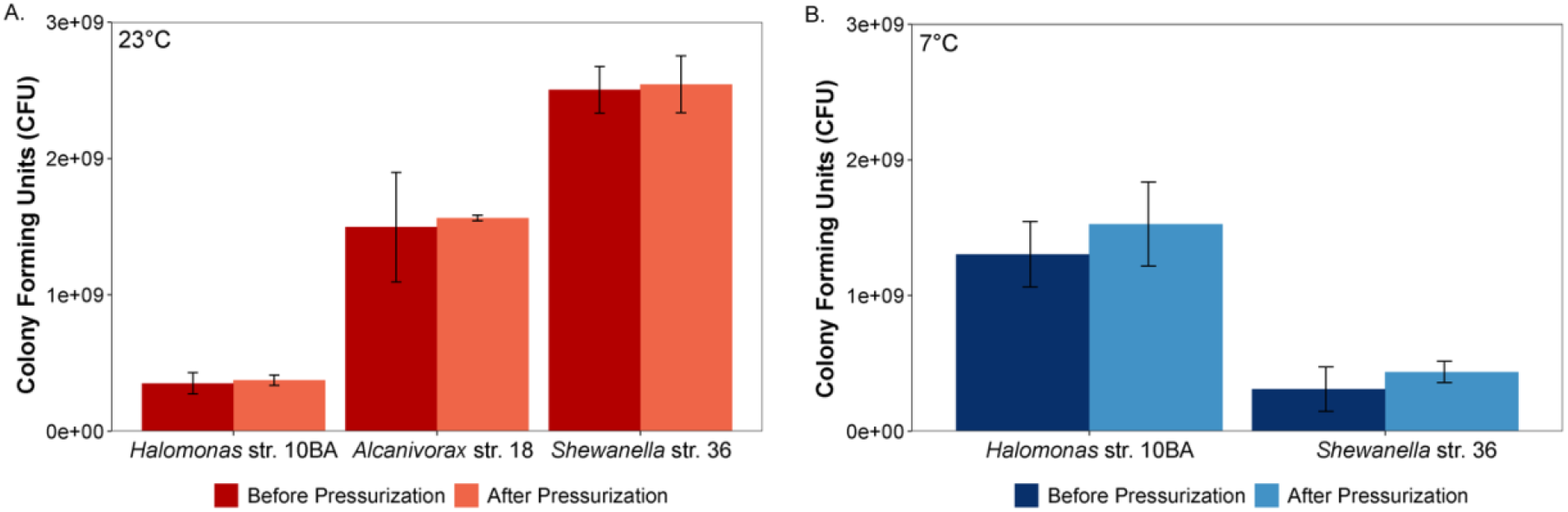
Colony forming units (CFU, n = 3) of the study strains before and after pressurization at (A) 23°C and (B) 7°C.

**Supplementary Figure S7.**
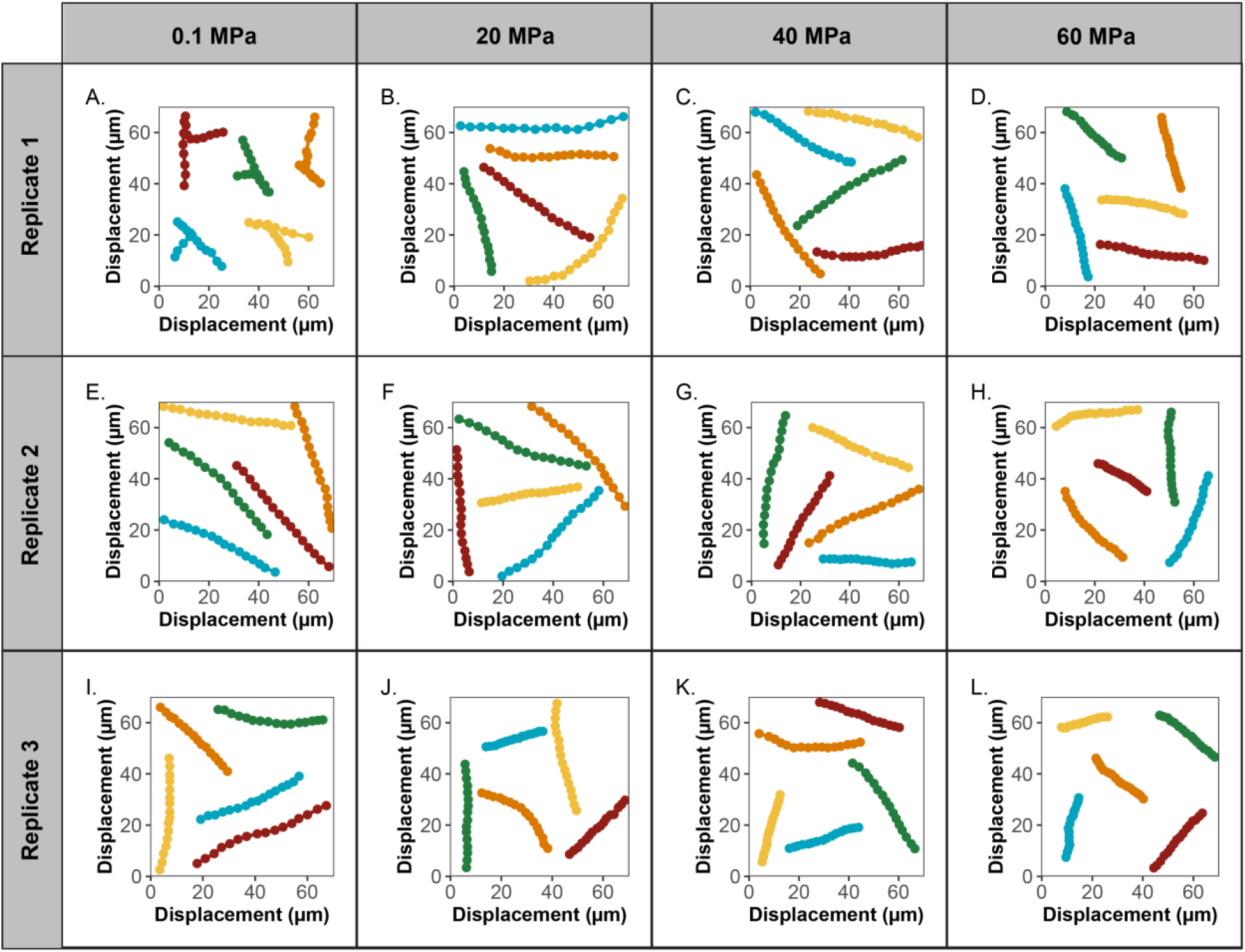
Trajectories of representative *Alcanivorax* strain 18 GOM-1409m cells at 23°C as a function of pressure. The positions of the cells were plotted every 2nd frame for 1 second. Trajectories for three replicates are shown (A-D are Replicate 1, E-H are Replicate 2, and I-L are Replicate 3). For each replicate, pressure increases from left to right.

## SUPPLEMENTARY VIDEOS

**Supplementary Video S1.** Compilation of high-pressure microscopy videos of *Shewanella* sp. strain 36 GOM-46m at 23°C. Pressure is increased in increments of 20 MPa over time to a maximum of 120 MPa, then is decreased in increments of 20 MPa over time (depressurization).

https://drive.google.com/file/d/1N4IaRJ2x3x7pYy5kngDtGA58OcOkvbW6/view?usp=sharing

**Supplementary Video S2**. High-pressure microscopy videos of *Alcanivorax* sp. strain 18 GOM-1509m Replicate 1 at 23°C. A clear decrease in switching frequency can be seen as pressure increases from 0.1 to 20 MPa. The first round of videos is at normal speed (30 frames/second), and the second round is slowed down 50% (15 frames/second).

https://drive.google.com/file/d/lcb8TvgRPPIXgbfe3zplKEvjx56jOviVo/view?usp=sharing

